# Beyond variability: a novel gene expression stability metric to unveil homeostasis and regulation

**DOI:** 10.1101/2024.05.28.596283

**Authors:** Mengjie Chen

**Author notes:** ***Corresponding authors*** Correspondence to Mengjie Chen.

## Abstract

The concept of gene expression stability within a homeostatic cell is explored through the gene homeostasis Z-index, a measure that highlights genes under active regulation in response to internal and external stimuli. This index reveals distinct regulatory activities and patterns in different organs, such as enhanced synaptic transmission in pancreatic islets. The research indicates that traditional mean-based methods may miss these nuances, underlining the significance of new metrics in identifying gene regulation specifics in cellular adaptation.

## Introduction

A homeostatic cell carries out regular functions to maintain balance within the cell. This involves continuously reacting to both internal and external stimuli, a process that often involves the regulation of transcription. Within a homeostatic cell, the majority of genes are transcriptionally stable, and a smaller proportion might be in a regulatory or compensatory state, acting as a response to stimuli. Key candidates for this type of regulation could be ‘first responder’ genes, interferons and genes that encode heat shock proteins, among many others. Here, we introduce the concept of gene expression stability, measured by the gene homeostasis Z-index. This index unveils genes subject to precise regulation within specific cell subsets, shedding light on their roles in cellular adaptation. For example, we observe organ-specific patterns exemplified by heightened synaptic transmission activities in islets. Furthermore, we uncover regulatory patterns for neuropeptides, such as insulin and somatostatin, exhibiting extreme values within a limited number of cells. These findings underscore the limitations of conventional mean-based approaches, highlighting our approach’s ability to surpass these constraints.

### K-proportion and gene homeostasis Z-Index

As cells of the same type and in the same microenvironment are likely to receive similar stimuli, leading to similar molecular profiles, we can leverage the statistical properties of cell clusters in single cell genomics data to differentiate between regulatory and homeostatic genes. To better distinguish between the gene regulatory states, we introduce a gene-specific statistic, ‘k-proportion’. This is the percentage of cells with expression level of a specific gene below a given integer value, k, which is determined based on the mean gene expression count (Figure 1, see Methods for details). Consider a situation where certain genes exhibit high expression levels in a limited subset of cells. This extreme expression skews the overall average upward, however the k-proportions (a measure reflecting the portion of cells with considerably lower expression) remain high. Essentially, under equivalent mean expressions, regulatory genes display significantly higher values of k-proportions compared to their homeostatic counterparts. This relationship between k-proportions and the mean expression can be illustrated on a ‘wave plot’. In this visualization, regulatory genes surface as anomalies, appearing as distinct outliers above the general trend, akin to water droplets cast airborne from a wave (Figure 1). By isolating these ‘droplets’, we can identify potential regulatory genes, offering new avenues for characterizing gene regulation directly from single-cell data. To quantify the extent of deviation from the homeostatic gene population, we propose a k-proportion inflation test that compares the observed k-proportion with the expected value from a set of negative binomial distributions with a shared dispersion parameter, which is estimated empirically assuming a majority of genes are homeostatic. Due to the asymptotic normality of the proposed test under the null hypothesis, we can obtain a Z-score for each gene, as a gene homeostasis Z-index, to serve as a measure of gene expression stability. Higher scores indicate more active regulation or compensatory activity.

**Figure 1.**
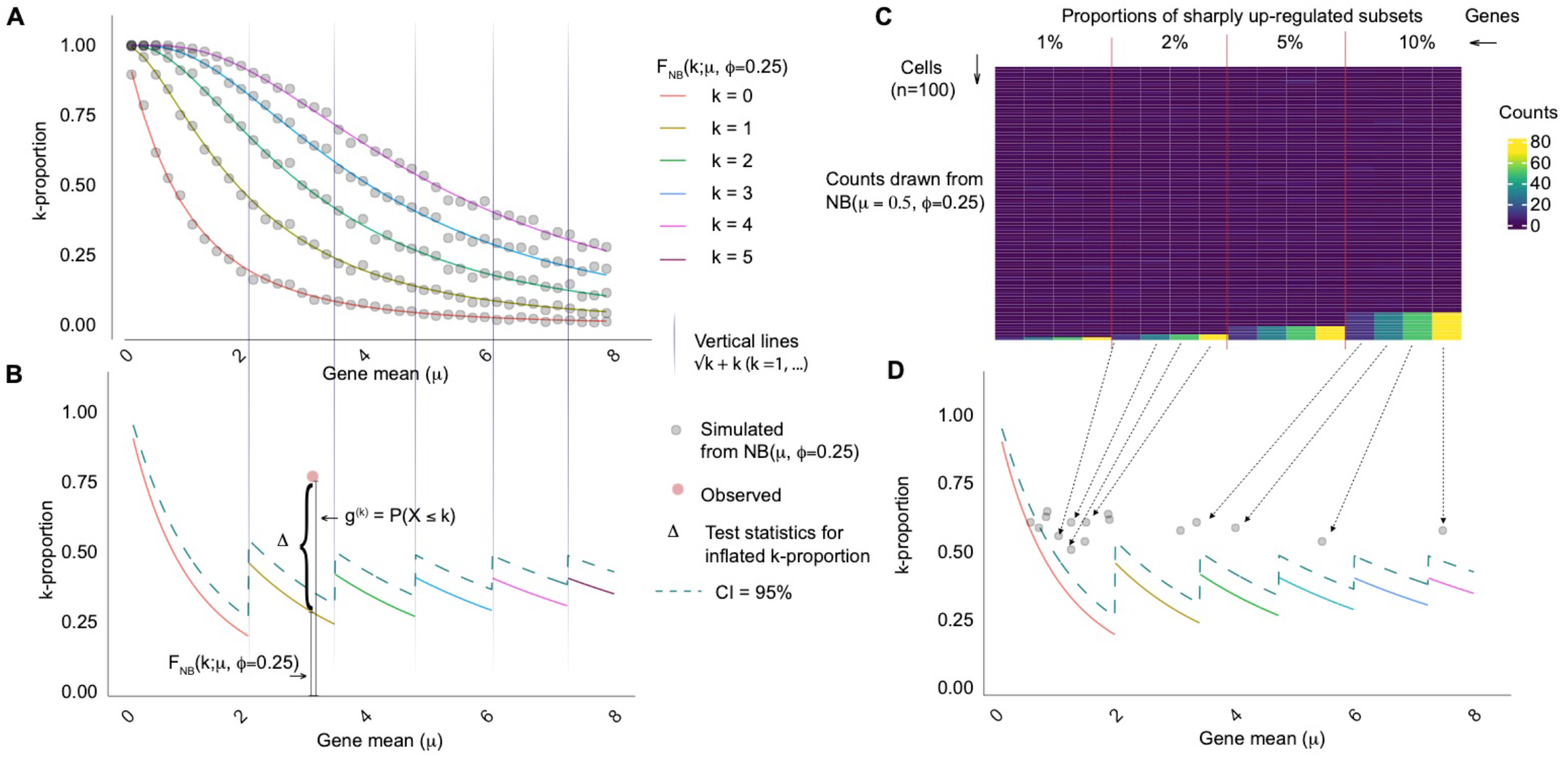
Illustration of k-proportion inflation test and the wave plot. k-proportion is defined as the proportion of counts no greater than k. In (A), we show k-proportions as a function of gene mean expected from a negative binomial distribution (curves) and from simulated data (dots), for k = 0, …, 4. For each gene, only one k is tested. The choice of k depends on the gene mean (vertical lines). In the test statistic 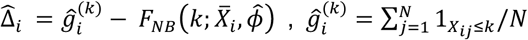, is essentially a nonparametric estimate of the true proportion, robust to the underlying distribution; 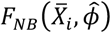 is a parametric estimate of the proportion, under the assumption of a negative binomial model. The test statistic compares the difference between a nonparametric and a parametric estimate (B, CI = confidence intervals). If the difference is significant, it suggests that negative binomial is not a good fit. Since we test for observed k-proportion being strictly larger, the rejected hypothesis indicates existence of highly expressed outliers, i.e., the gene expression in up-regulated in a subset. In a wave plot, where k-proportion is displayed as a function of gene mean, regulatory genes surface as anomalies, appearing as distinct outliers above the general trend (C, D). The gene homeostasis Z-index is the z-score derived from the test on k-proportion.

We examined the distribution of k-proportions in a dataset of CD34+ cells (Figure 2)^1^. Through UMAP visualization, we identified three distinct subgroups within the CD34+ cell population. We generated wave plots for each subgroup separately and for all subgroups combined. Consequently, we observed that the empirical distribution of k-proportions in homeostatic genes (most genes) closely aligns with those from negative binomial distributions with the same dispersion level. In relatively homogeneous cell populations, the resulting distribution approximates those from a Poisson distribution. In subgroup 3, the dispersion level is 0.163, indicating similarity to a Poisson. However, as cellular heterogeneity increases, so does the dispersion level. In subgroup 2, the dispersion level is 0.526, indicating increased variability in gene expression. When considering all subgroups together, the dispersion level is 1.4. In all these scenarios, we consistently observe ‘droplets,’ which signify genes undergoing active regulation. Figure 2(CD) displays selected genes with significant Z-index after adjusting for multiple comparisons using FDR, for each cell population (Figure S1). In subgroup 1, H3F3B, GSTO1, TSC22D1, CLIC1, LYL1 and FAM110A are under regulation in a subset of cells, indicating cellular oxidant detoxification activities. In subgroup 2, PRSS1 and PRSS3 are upregulated a small portion of cells, revealing digestive activities. In subgroup 3, NKG7 and GNLY are upregulated, for cell-killing activities. Across all combined groups, we observe active regulation of HLA and RPL families, exclusively occurring in subgroup 2, indicating cytoplasmic translation and processing of exogenous peptide antigen activities. Additionally, MAP3K7CL is upregulated in subgroup 1, suggesting this cell population might transmit signals downstream in response to extracellular stimuli through MAPK signaling pathway. Overall, we show Z-index can be applied to both homogeneous and heterogenous cell groups to identify sharply up-regulated genes.

**Figure 2.**
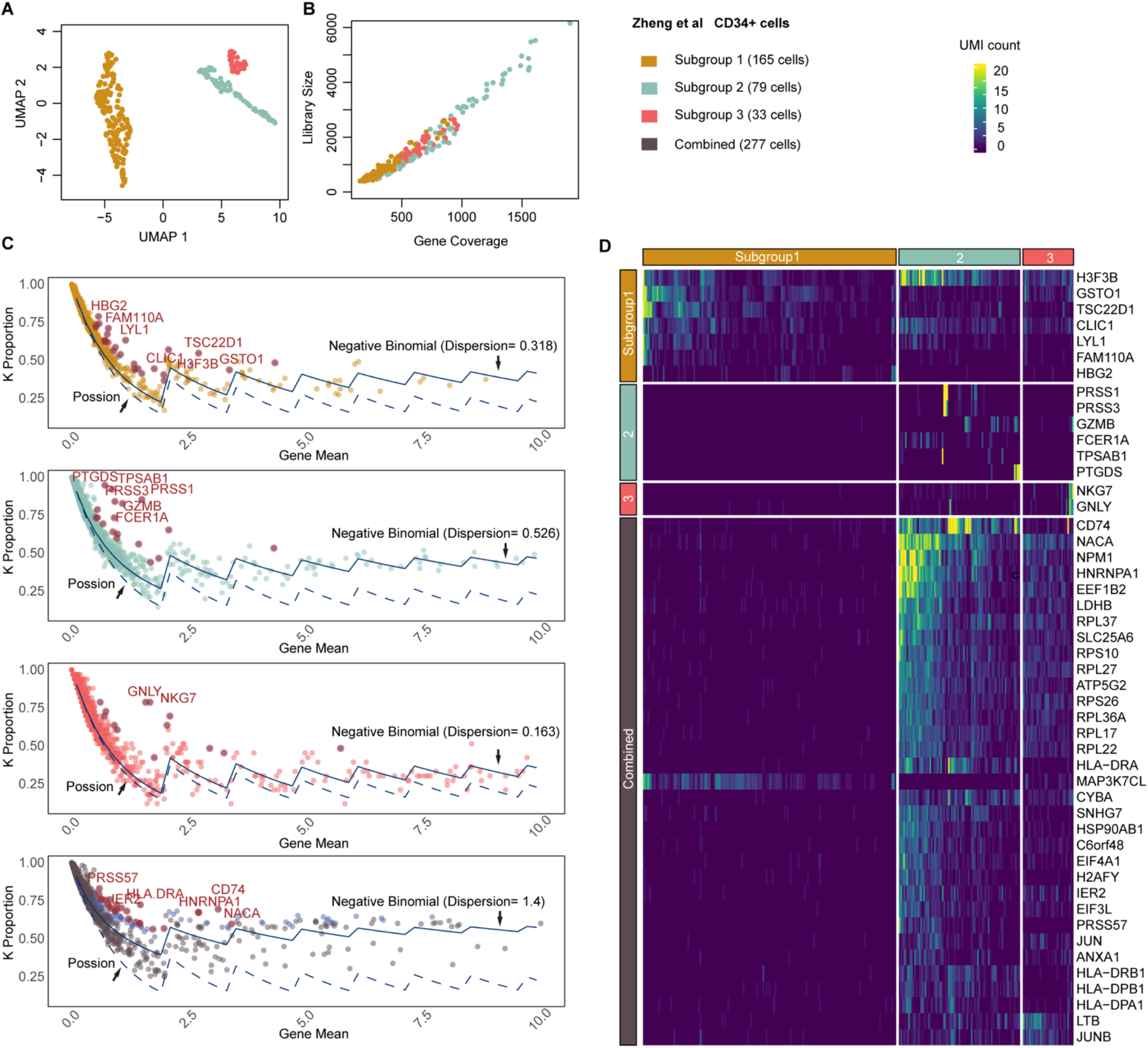
Gene expression stability analysis on Zheng et al CD34+ cells. A. UMAP show three subgroups in the CD34+ cells. B. Library size vs gene coverage for each individual cell. C. Wave plots of observed k-proportion vs gene mean for subgroup 1, 2, 3 and all subgroups combined. Genes with top significant Z-index are labeled by their names. The k-proportion vs gene mean from a Poisson distribution and from the fitted negative binomial are shown. D. Heatmap visualization of selected regulatory genes for cells from different subgroups.

### Gene expression stability captures information beyond variability

Gene expression variability quantifies the degree of fluctuation in a gene’s expression levels relative to its mean expression value. This is commonly measured using statistical parameters such as variance or coefficient of variation (CV)^2-8^. These metrics are often used in gene feature selection, as genes with higher variability are believed to provide more information due to their pronounced expression fluctuations across diverse cells. Contrastingly, gene expression stability provides an alternative perspective. It centers on a gene’s baseline expression status and its consistency in maintaining this status across different cells. Genes with lower stability are more likely to undergo sharp regulation within specific cell subsets. Importantly, genes with lower stability can exhibit either low or high variability, highlighting the diverse aspects of gene expression that these metrics capture. We demonstrate the differences between these metrics through 1,011 cells from a β cell subtype from an individual donor. After correcting for multiple comparisons, we identified 372 genes out of 18,513 coding genes that have significant hemostasis Z-indexes and with k-proportions inflated over 10%. In genes like SYT16, RBP4, PCDH7 and IAPP, ‘droplets’ show leading inflated k-proportions on a wave plot and exhibit moderate variance and CV^2^ (Figure 3). Interestingly, genes with lower stability can have higher variance under the same mean expression level, while genes with high variability can remain relatively stable across the entire cell population. Among the genes with top variance but relative stability is LRRTM4, G6PC2, NRG3, CNTN4 and CNTN5 (Fig S2). We also investigate the relationship between Z-index, variance, and CV. Interestingly, we find no monotonic relationship across all genes, indicating that these metrics depict different properties. However, for genes with a fixed mean expression, those with a high Z-index exhibit greater variance. Upon a detailed examination of the expression patterns and distributions of the top inflated genes across cells (Figure 3C-D), we observe significant variations in both the proportions and expression levels of up-regulated genes. This observation underscores the flexibility of the Z-index, as it can capture a broad spectrum of expression patterns originating from localized up-regulation, without any prerequisite assumption of the underlying distribution.

**Figure 3.**
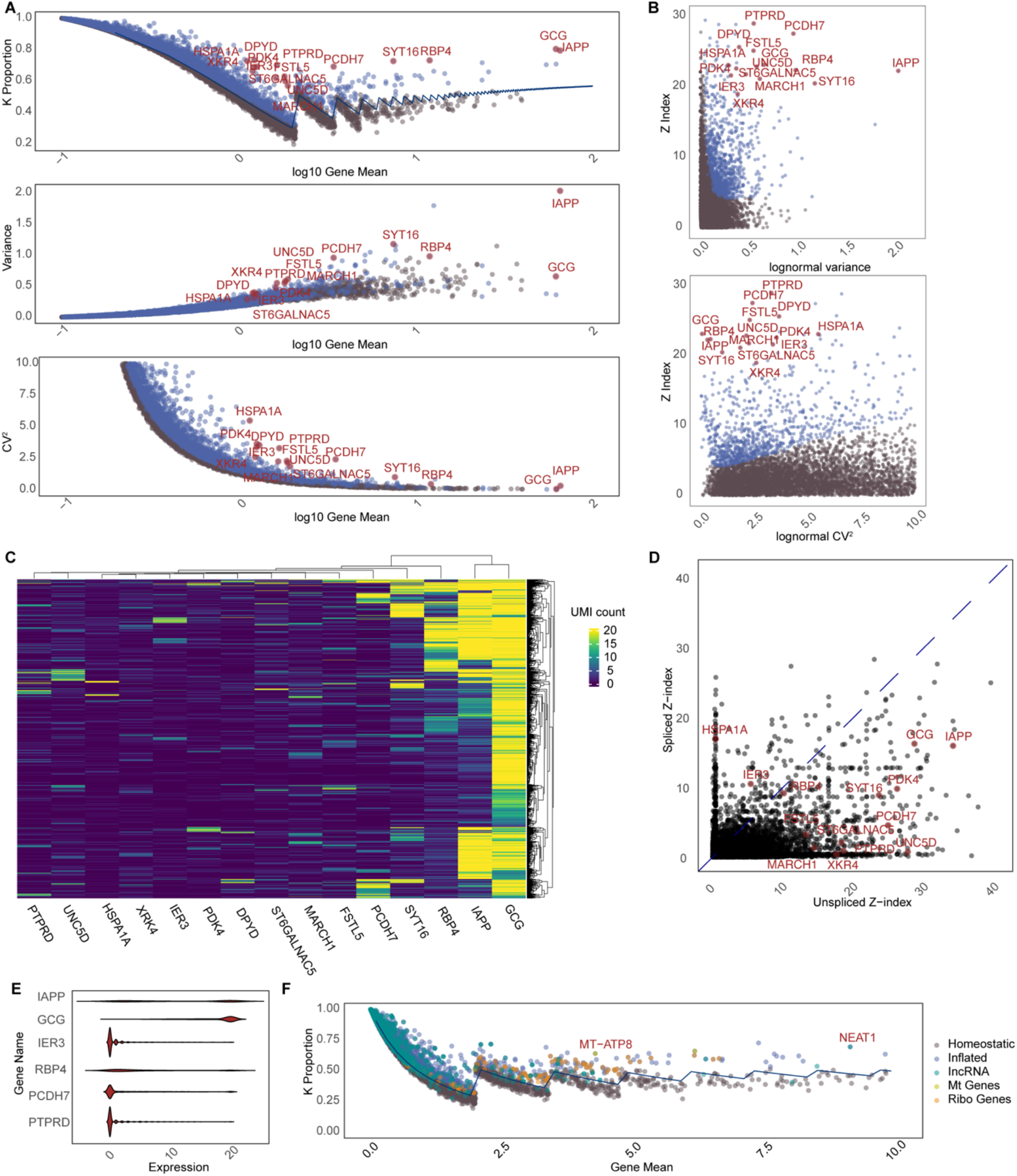
Gene expression stability analysis on pancreatic β cells from one donor. Top 15 genes from the inflation test are denoted in red in all plots. A. K-proportion, variance and CV^2^ as a function of gene mean (log 10 scale). B. Scatterplots of Z-index vs. variance and Z-index vs. CV^2^. C. Heatmap of UMI counts for top 15 inflated genes across all cells. D. Z-indexes derived from separate tests on spliced UMI count and un-spliced UMI counts. E. Violin plot shows heterogeneity of count distribution across top 15 inflated genes. F. Wave plot shows k-proportions derived from long non-coding RNAs (lincRNA), mitochondrial genes’ RNAs, and ribosomal genes’ RNAs, which all can be fitted with the same negative binomial distribution as the coding genes.

Velocity analysis has gained popularity for studying gene expression dynamics by distinguishing between nascent (spliced) and mature (unspliced) RNA molecules^9^. This inspired us to assess the stability Z-index for spliced and unspliced counts, respectively. Intriguingly, the k-proportions for these counts each adhere to a distinct group of negative binomial distributions, both maintaining the same dispersion level (Fig S3). Furthermore, we find that up-regulation of gene expression can be achieved through different avenues: some genes amplify their spliced counts, others their unspliced counts, and yet others both. For instance, among the top 15 genes, HSPA1A and IER3 only show inflated spliced counts in specific cell subsets. Conversely, UNC5C, UNC5D, and MARCH1 primarily demonstrated an increase in un-spliced counts. Genes like GCG, IAPP, SYT16, RBP4, and PDK4 exhibited elevations across the board. These findings emphasize the nuanced nature of gene expression up-regulation, suggesting influences beyond mere transcriptional boosts, possibly involving independent mechanisms like accelerated splicing or reduced degradation.

It is noteworthy that k-proportions derived from long non-coding RNAs (lincRNA), mitochondrial genes’ RNAs, and ribosomal genes’ RNAs all can be fitted with the same set of negative binomial distributions as the coding genes in the nucleus (Figure 3F). The shared distribution may hint at underlying biological or biochemical processes that govern RNA production or stability across these diverse RNA types.

### Gene homeostasis Z-index as a portable self-normalizing metric

Gene homeostasis Z-index provides a measure of gene expression stability across genes within a cell group. It inherently absorbs both technical and biological variability within the dispersion term of the negative binomial distribution. As such, the Z-index alleviates concerns related to batch effects and other unwanted influences. This feature makes the Z-index a portable self-normalizing metric for comparisons across individuals and datasets. We applied the gene homeostasis Z-index to a dataset from pancreatic islets, sourced from 29 individual donors with Type 2 diabetes (T2D) or Non-diabetic (ND). Insulin secretion is a tightly regulated process that responds rapidly to changes in glucose concentration. In the context of T2D, one central issue is impaired insulin secretion and function. This makes T2D an ideal disease to demonstrate the effects of the Z-index and its relevance to the disease state. Our analysis extended to each of the 15 cell types documented in the original study, evaluated individually for every donor. Consequently, the derived Z-index gauges the stability of a gene within a particular cell type from a specific donor.

From our results, we identified 1,617 genes that displayed significant up-regulation in a subset of cells, after adjusting for the FDR. Among these, some are generic genes that respond to cell stress, including immediate responders, JUN and IER3, heat shock responders and inflammatory genes. GO analysis revealed that these regulatory genes are enriched in metabolic functions, such as hormone metabolic process and peptide hormone secretion (Figure 4).

**Figure 4.**
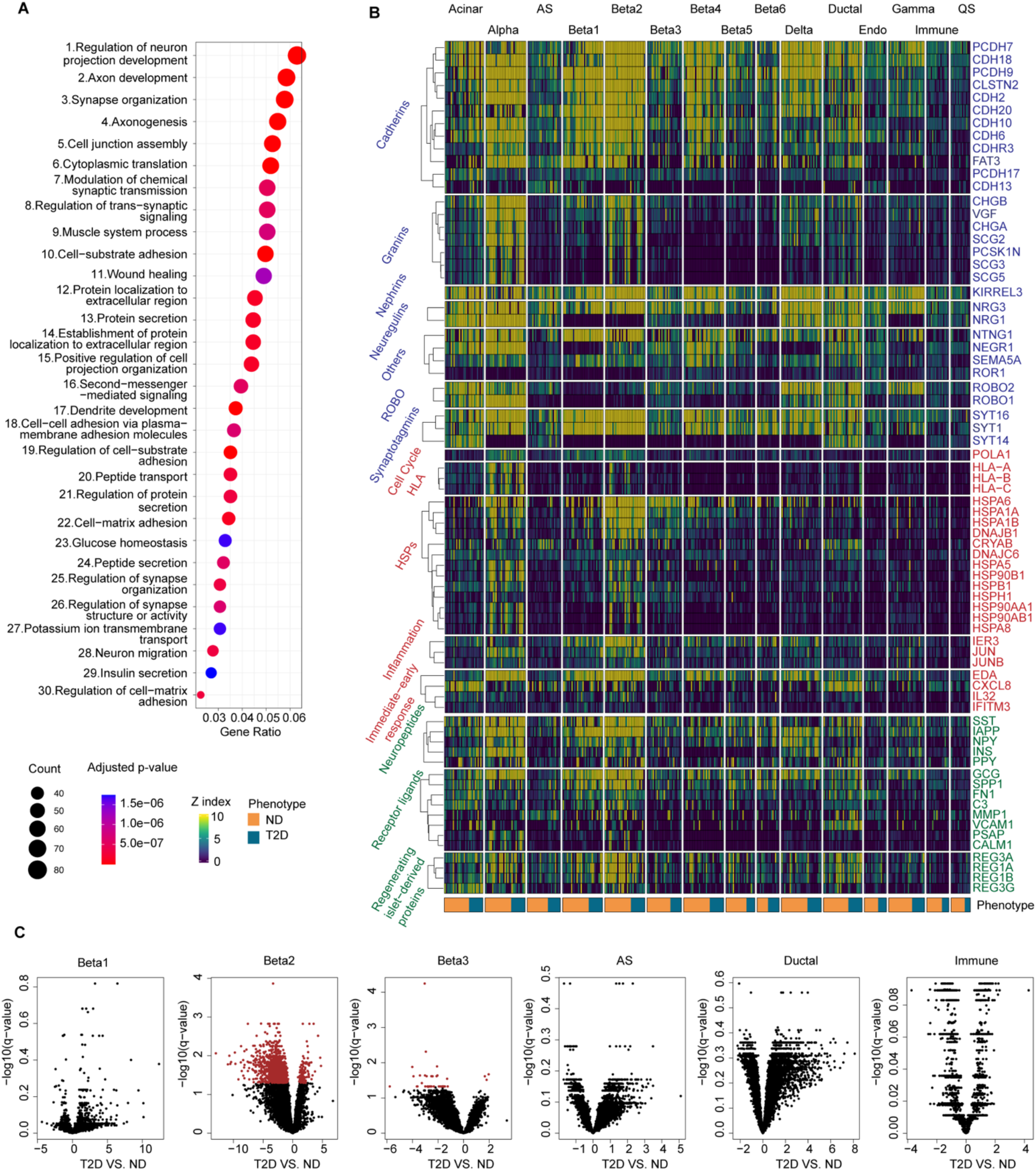
Gene expression stability analysis on pancreatic cells from multiple donors. A. Top 30 enriched GO categories from GO enrichment analysis on 1,617 genes that displayed significant up-regulation. B. Heatmap visualization of Z-index from selected regulatory genes across different cell types across individual donors. Genes are organized into three categories: neuronal genes (blue), generic first responders (red), islet secretion related genes (green). C. Volcano plots of differential stability analysis between T2D and ND across cell types.

Notably, we also find significant enrichment of neuronal functions in islet cell types, including various synaptic transmission and dendrite activities. It is known that both the central nervous system and autonomic nervous system regulate and impact the pancreas for insulin secretion and glucose control. However, the characterization of a hypoglycemia sensing and counter-regulatory system within the pancreas is still in its infancy. Our analysis highlighted consistent regulatory activities of genes like cadherins, granins, nephrins, neuregulins, ROBO, and synaptotagmins across varied islet cell types and individuals. While the overall pattern of the Z-index for selected neuronal markers in α, β_1_, β_2_, δ and Ductal cells are similar, we observe stronger ROBO1, NRG1 and NEGR1 up-regulation in α cells, SYT1 and NTNG1 up-regulation in β_1_ and PCDH7, KIRREL3, NRG3 and SYT16 up-regulation in β_2_, hinting at cell-specific inclinations even within gene families (Figure 4). The difference of regulatory activities cannot be revealed by mean-based approaches (Figure S4).

We then performed differential stability analysis within each cell type, by comparing T2D with ND cases. We observed significant differential stability primarily in β cells, specifically β_2_ cells. Other cell types did not display significant differences. Notably, genes in the T2D-affected β cells showed a tendency for decreased up-regulation, as indicated by smaller Z-index values compared to ND cases. Many of these genes are linked to neuronal activities, including PDE4B, NCAM2, SEMA3A, DPP10, CHODL, UNC5C, PTH2R and EPB41L4A, constituting a list of 116 genes with Z-index differences greater than 5. To identify key master regulators of these 116 genes, we employed ChEA3^10^, a transcript factor enrichment analysis tool that integrates multiple omics data. The top 10 transcription factors identified were NPAS3, ZNF385B, DBX2, ZNF385D, MEIS2, RORB, PURG, ZNF804A, TBX18, and PRRX1. Notably, NPAS3, DBX2, and MEIS2 are known to regulate genes involved in neural development^11-13^. Importantly, chromosomal abnormalities affecting the coding potential of these genes have been associated with conditions such as schizophrenia, cognitive disability, and obesity. Notably, few of these TFs are documented in JASPER database. They can be overlooked if using a motif-based approach, even with paired ATAC-seq data. For β_2_ cells, we performed differential expression analysis using gene mean for T2D vs. ND and compared log fold-change vs. differences in Z-index. We found differential stability analysis indeed pinpointed to a different gene set, the scope of which was different from mean-based approaches (Figure S5).

Furthermore, we investigated whether the up-regulation of genes occurred in the same group of cells by assessing their correlations. Our results on β_2_ cells across all donors indicated a strong correlation in the expression levels of neuregulins, ROBO, and ROR1. Another gene module encompassed genes such as INS, IAPP, HSP90, HLAs, and granins. A separate cluster was dominated by JUN and several HSPs. Furthermore, REGs and IL32 collectively constituted another highly correlated gene module. Such patterns indicate that individual β_2_ cell subsets may assume specialized roles influenced by their unique microenvironments. These roles range from interfacing with the nervous system and producing insulin to promoting cell proliferation, with some subsets potentially multitasking across several roles (Figure 5).

**Figure 5.**
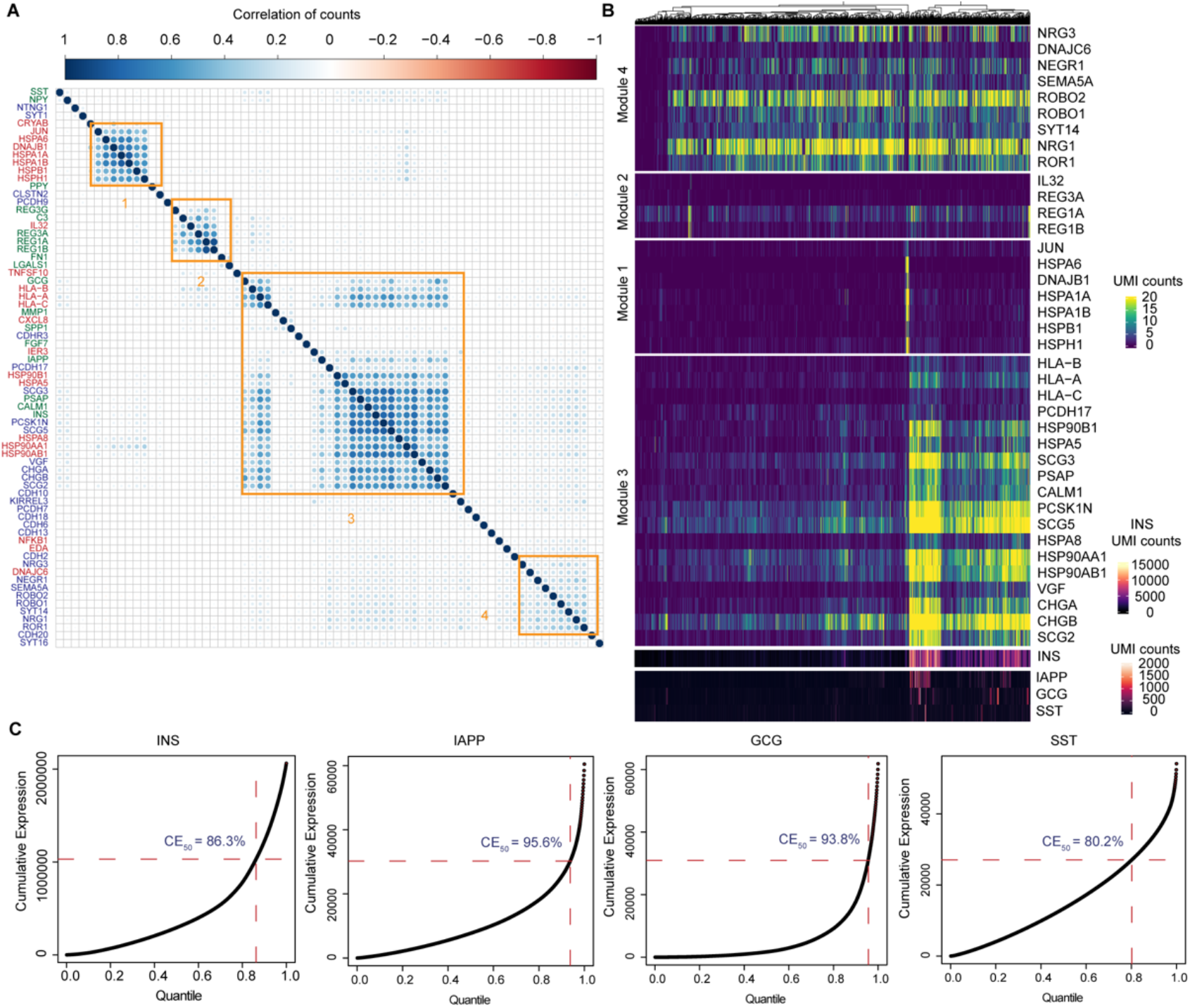
Further characterization of selected regulatory genes in β cells from one donor. A. Correlation of UMI counts of selected genes across all cells. Genes are grouped into modules by their correlations (orange squares). B. Heatmap visualization of gene modules identified from A. C. Example genes that exhibit extreme values. This deviation from a homogeneous distribution underscores a potential flaw in the current analytical methodologies that heavily rely on mean values.

Lastly, INS expression is notably up-regulated, albeit within a specific subset of cells, in tandem with the up-regulation of HSP90s and granins. Instead of following a negative binomial distribution, the INS expression pattern exhibits numerous extreme values. To illustrate, 13.7% of cells account for 50% of the total INS expression, while the final 5% are responsible for 25% of the overall expression (Figure 5). This deviation underscores a potential flaw in the current analytical methodologies that heavily rely on mean values. Our observation suggests there may be a baseline expression of INS across beta cells—indicating a latent potential for insulin production. The rapid synthesis in response to external stimuli, such as glucose levels or nervous system signals, seems to be a specialized function of cell subsets. The regulatory patterns observed for other important endocrine factors, IAPP, SST and GCG, mirror this principle, where localized extreme values significantly contribute to their overall high expression in β cells.

## Discussion

In the field of single-cell research, the focus has primarily centered around studying the characteristics of cell types with uniform and consistent behaviors. Consequently, genes that exhibit more varied and inconsistent behaviors have often been overlooked^14-16^. However, these very genes, tightly regulated in specific cell subsets, hold immense importance as they enable cells to respond and adapt to diverse internal and external stimuli within their microenvironment. Recognizing the fundamental role of these genes in maintaining cellular homeostasis, we introduced a new concept - gene expression stability.

Gene expression stability focuses on the consistent baseline expression status of a gene across different cells in a sample, providing insights into which genes undergo sharp regulation within specific subsets. The gene homeostasis Z-index serves as a quantitative measure of gene expression stability. High scores indicate active regulation or compensatory activity, shedding light on which genes are actively regulated in response to stimuli. Using the Z-index, we observed significant regulatory undertakings in generic defense mechanisms, such as cellular stress response and inflammation. Additionally, organ-specific activities were observed, notably synaptic transmission activities in islets. Crucially, we observe the regulatory patterns for important neuropeptides, including INS, IAPP, and SST, are highly localized, showing extreme values in a limited number of cells. This observation emphasizes the shortcomings of current methods that predominantly consider mean expression.

Our study points to the importance of β cells and their reduced neural gene regulation in T2D. Understanding these specific genetic and regulatory changes could lead to more targeted and effective treatments for T2D. This might involve interventions aimed at modulating the activity of master regulators like NPAS3, DBX2, and MEIS2. Additionally, we anticipate that the Z-index could advance research into conditions characterized by impaired stimulus response or metabolic dysfunction, such as obesity and cardiovascular diseases.

Previous methodologies have attempted to fit curves between CV and the mean, assuming that technical effects can be encapsulated by this general trend^2, 17, 18^. These models then pinpoint genes deviating from this trend as noteworthy. While this strategy holds merit for feature selection, it faces challenges of controlling false discoveries. Moreover, it neither produces transferable statistics nor directly elucidates gene regulation mechanisms. In contrast, the gene homeostasis Z-index offers a detailed snapshot of gene expression stability within a single sample, across RNA types and spliced/unspliced transcripts. It excels as a portable metric, offering consistent, comparative insights across multiple samples. This portability, combined with its ability to capture nuanced expression dynamics, makes the Z-index a promising tool in single-cell genomics and broader transcriptomic analyses.

## Methods and Materials

### *k*-proportion and the *k*-proportion inflation test

Denote the number of cells as *N*, the number of genes as *M*, and let the observed gene expression matrix (UMI counts) be denoted as *X*. Each element indexed by *X*_*ij*_ represents the UMI count of the *i*-th gene in the *j*-th cell. We define *k*-proportion as the percentage of cells with gene expression less than a defined integer value 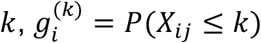. Our goal is to test whether the underlying true *k*-proportion is significantly higher than the expectation under a negative binomial model, i.e.,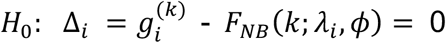, where *F*_*NB*_ is the cumulative distribution function under a negative binomial distribution, *λ*_*i*_ is the mean and *ϕ* is the dispersion level, i.e., *F*_*NB*_ (*k*) = *P*_*NB*_ (*X*_*ij*_ ≤ *k*). Under the alternative hypothesis, *k*-proportion is inflated due to active regulation in a small subset, i.e., 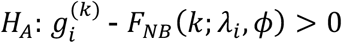. Since 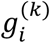, mean *λ*_*i*_ and dispersion level *ϕ* are unknown, we replace these parameters by their estimates, respectively. Observed *k*-proportion for *i* -th gene for a given *k*, is computed as 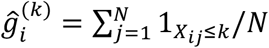. The observed mean expression is estimated as 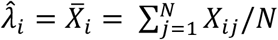. Our test statistic compares the difference between observed *k*-proportion and the fitted value under a negative binomial model: 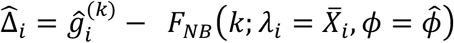. The estimate of *ϕ* and choice of *k* will be discussed as follows.

### Estimation of dispersion level

Within the same cell type from a single cell experiment, we observed that the empirical distribution of *k* -proportions in most genes (homeostatic) closely aligns with *k* - proportions that are generated from negative binomial distributions with the same dispersion level across genes. Since the dispersion level captures the variability introduced by technical factors and biological factors, the observation confirms that these factors affect cells measured in the same single cell experiment the same way. We estimated the dispersion level by minimizing the square error loss between fitted *k* -proportions and observed values across all genes: 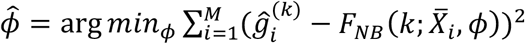. We assume homeostatic genes outnumber regulatory genes by a magnitude; thus, no filtering on genes was performed.

### Choice of *k*

Since we use *k*-proportion as a measure reflecting the portion of cells with considerably lower expression, it is easy to see that for a given mean expression, there can be multiple choice of *k*. Intuitively, we can compute any proportion of values smaller than the gene mean. However, if *k* is chosen too small, it cannot effectively discern the differences. If *k* is chosen too large, it cannot well represent low expression. Furthermore, we can choose only one *k* for each gene to make sure all tested hypotheses are independent so that the false discovery control is straightforward. For simplicity, setting *ϕ* = 0, we choose the smallest integer value that density *f*_*NB*_(*k*; *λ*_*i*_, 0) is monotonically decreasing given 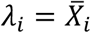. We can show that such *k* can be obtained by satisfying 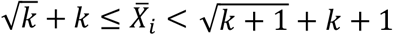.

### Gene Homeostasis Z-index

After determining *k* and 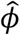, our test statistic is the difference between observed and expected *k*-proportion: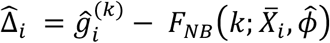. The detection of regulatory genes is now equivalent to a one-tailed test against the null, Δ_*i*_ = 0. Under the alternative, Δ_*i*_ > 0. A Z-index (Z-score) can be computed for each gene as 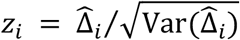. We can calculate the variance of Δ_*i*_ using the delta method, which leads to the following equation:

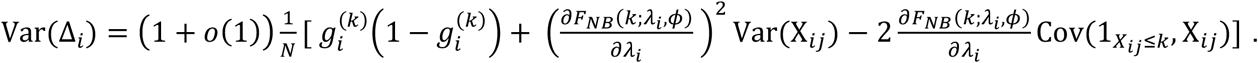

The first term accounts for the variability of 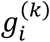, the second term accounts for the variability of *F*_*NB*_(*k*; *λ*_*i*_, *ϕ*) and the last term is the covariance between 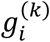 and *F*_*NB*_(*k*; *λ*_*i*_, *ϕ*). We compute Var 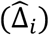 by plugging in estimates under the null, i.e., 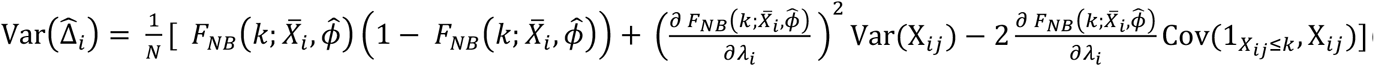 (See Appendix for details). A *p*-value can be derived from the Z-score using the standard normal distribution. We used Benjamini Hochberg method to adjust for multiple comparisons across genes.

### Interpretation

In the test statistic 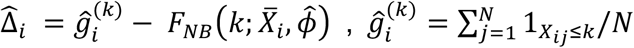 is essentially a nonparametric estimate of the true proportion, which is robust to the underlying distribution; 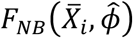 is a parametric estimate of the proportion, under the assumption of a negative binomial model. Our test statistic compares the difference between a nonparametric and a parametric estimate to see whether there is significant discrepancy between the two. If the difference is significant, it suggests that negative binomial is not a good fit. Since we test for observed *k*-proportion being strictly larger, the rejected hypothesis indicates existence of highly expressed outliers.

### Simulations

We simulate data under the null hypothesis, examining various scenarios with different means and dispersion levels. For each scenario, we generate 1000 independent datasets and subsequently derive z-scores using the k-proportion inflation test. Our observations reveal that the distribution of these z-scores closely approximates a normal distribution with a mean of 0 and a standard deviation of 1. This observation indicates that the z-score distribution adheres to a normal distribution asymptotically. Consequently, we find that the type I errors for these tests are well-calibrated (Figure S6).

We next assess the effectiveness of the proposed test, taking into account various factors that can influence its power. These factors encompass dispersion levels, cell numbers, the proportion of genes with sharp regulation, and their mean expression. Specifically, we fix the cell number at 200. Within this framework, we perform simulations where we generate baseline expression data following a negative binomial distribution with dispersion parameters of 0.5 (empirical estimates based on real data). Additionally, we introduce up-regulated expression data with fixed values of 2, 4, 8, and 10. The percentage of genes exhibiting up-regulation varies across simulations, including scenarios with 1%, 2%, 5%, and 10% of genes showing increased expression (Figure S7-8). In brief, for lowly expressed genes (gene mean = 0.1), we can achieve a power of 75% for 1% of outlier with mean greater than 4. For higher expressed genes, to reach a decent power, we need to increase either the percentage of outliers or the expression level of outliers. In all cases, type I errors are well-controlled.

### Datasets

Throughout the analysis, we used publicly available single cell sequencing data from the 10X protocols. All the data sets were analyzed after their own filtering process. No extra filtering was performed.

Zheng et al data was downloaded from the 10X website. Only cells that labeled as CD34+ were selected for analysis. There are 277 cells with 31482 genes. We performed UMAP on log-transformed data. We used k-means to separate cells into 3 subtypes using their UMAP coordinates. We applied the k-proportion inflation test on UMI counts for each subtype and with all subtypes combined.

T2D data is downloadable from GSE234313. There are 160,954 cells with 32,746 genes, from 29 donors. In the original dataset, cells are annotated into 15 cell types/subpopulations. For each gene in each cell, we have UMI counts for all transcripts, spliced and unspliced. We extended the k-proportion inflation analysis to each of the 15 cell types/subpopulations.

### Downstream analysis

The GO enrichment analysis was performed using ClusterProfiler. Transcription factor enrichment analysis was using ChEA3. The visualization was performed using ComplexHeatmap.

## Code availability

We provide an R package that implements proposed methods discussed in this study. The RegulationIndex package is available from GitHub (https://github.com/ChenMengjie/RegulationIndex). In addition, the R source code to reproduce all data analysis in the study is available from GitHub at https://github.com/ChenMengjie/RegulationIndex_Paper_2024.

## Acknowledgements

The work was supported by National Institutes of Health grant R01 GM126553, R01 HG011883 and HG012927, and additional grant no. NSF 2016307 and Sloan Research Fellowship to M.C.

## Author information

### Contributions

M.C. conceived this work, developed the methods, performed the analyses and wrote the paper.

## Ethics declarations

Ethics approval is not applicable to this study.

## References

1. Zheng, G.X. et al. Massively parallel digital transcriptional profiling of single cells. Nature communications 8, 14049 (2017).

2. Arzalluz-Luque, Á., Devailly, G., Mantsoki, A. & Joshi, A. Delineating biological and technical variance in single cell expression data. The international journal of biochemistry & cell biology 90, 161–166 (2017).

3. Eling, N., Richard, A.C., Richardson, S., Marioni, J.C. & Vallejos, C.A. Correcting the mean-variance dependency for differential variability testing using single-cell RNA sequencing data. Cell systems 7, 284-294. e212 (2018).

4. Hafemeister, C. & Satija, R. Normalization and variance stabilization of single-cell RNA-seq data using regularized negative binomial regression. Genome biology 20, 296 (2019).

5. Jiang, H., Sohn, L.L., Huang, H. & Chen, L. Single cell clustering based on cell-pair differentiability correlation and variance analysis. Bioinformatics 34, 3684–3694 (2018).

6. Lun, A.T., McCarthy, D.J. & Marioni, J.C. A step-by-step workflow for low-level analysis of single-cell RNA-seq data with Bioconductor. F1000Research 5 (2016).

7. Buettner, F. et al. Computational analysis of cell-to-cell heterogeneity in single-cell RNA-sequencing data reveals hidden subpopulations of cells. Nature biotechnology 33, 155–160 (2015).

8. Huang, M. et al. SAVER: gene expression recovery for single-cell RNA sequencing. Nature methods 15, 539–542 (2018).

9. La Manno, G. et al. RNA velocity of single cells. Nature 560, 494–498 (2018).

10. Keenan, A.B. et al. ChEA3: transcription factor enrichment analysis by orthogonal omics integration. Nucleic acids research 47, W212–W224 (2019).

11. Lupo, G. et al. Molecular profiling of aged neural progenitors identifies Dbx2 as a candidate regulator of age-associated neurogenic decline. Aging Cell 17, e12745 (2018).

12. Pickard, B.S., Pieper, A.A., Porteous, D.J., Blackwood, D.H. & Muir, W.J. The NPAS3 gene—emerging evidence for a role in psychiatric illness. Annals of medicine 38, 439–448 (2006).

13. Machon, O., Masek, J., Machonova, O., Krauss, S. & Kozmik, Z. Meis2 is essential for cranial and cardiac neural crest development. BMC developmental biology 15, 1–16 (2015).

14. Jiang, L., Chen, H., Pinello, L. & Yuan, G.-C. GiniClust: detecting rare cell types from single-cell gene expression data with Gini index. Genome biology 17, 1–13 (2016).

15. Jindal, A., Gupta, P. Jayadeva & Sengupta, D. Discovery of rare cells from voluminous single cell expression data. Nature communications 9, 4719 (2018).

16. Liang, S. et al. Single-cell manifold-preserving feature selection for detecting rare cell populations. Nature computational science 1, 374–384 (2021).

17. Brennecke, P. et al. Accounting for technical noise in single-cell RNA-seq experiments. Nature methods 10, 1093–1095 (2013).

18. Stegle, O., Teichmann, S.A. & Marioni, J.C. Computational and analytical challenges in single-cell transcriptomics. Nature Reviews Genetics 16, 133–145 (2015).

