## Supplementary files for "Beyond variability: a novel gene expression stability metric to unveil homeostasis and regulation"

### Supplementary Figures

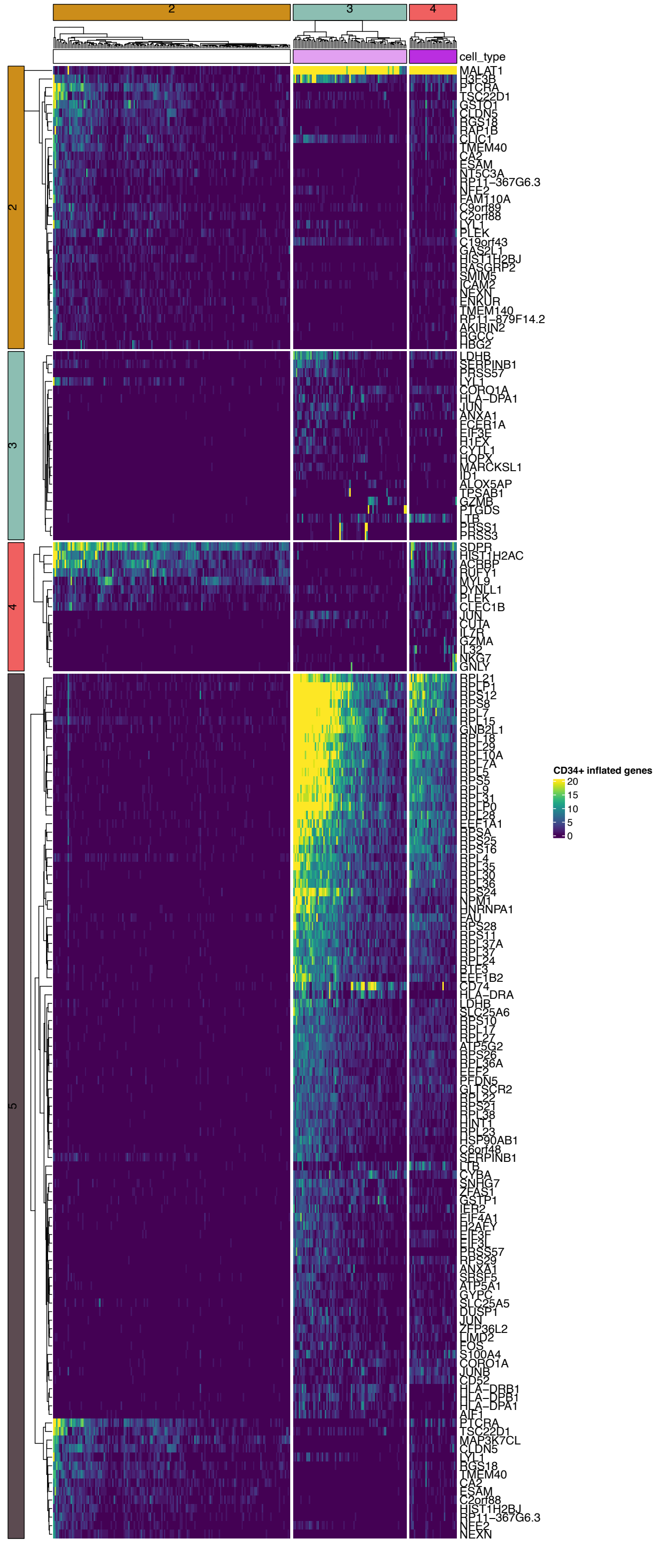


**Figure S1. Heatmap displays all genes with significant Z-index after adjusting for multiple comparisons using FDR, for each cell population from Zheng et al CD34+ cells.**

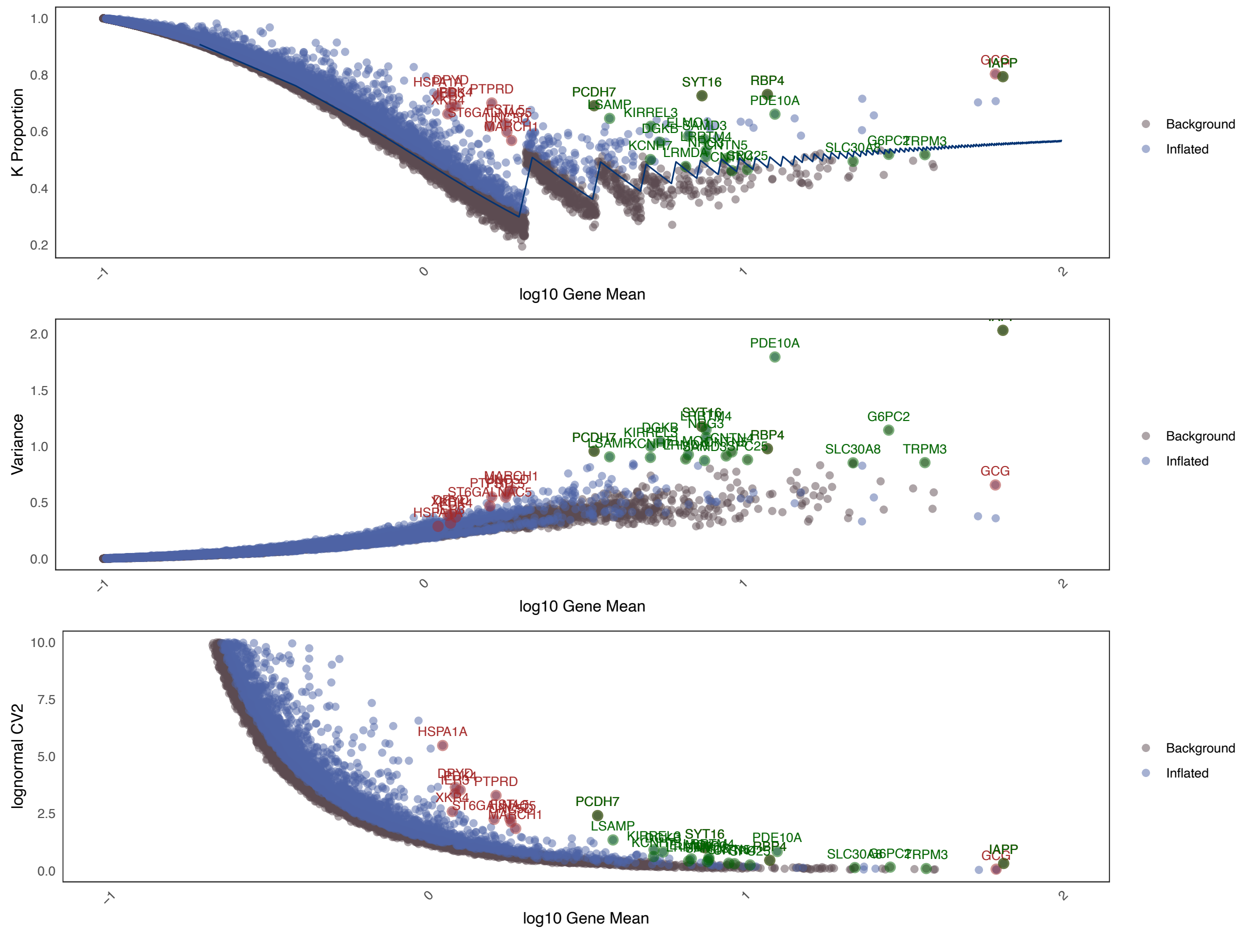


**Figure S2. Gene expression stability analysis in beta cells from the islet of one donor. Top varied and top inflated are shown in green. Top inflated but not top varied are shown in red. As shown, genes with lower stability can have higher variance under the same mean expression level, while genes with high variability can remain relatively stable across the entire cell population.**

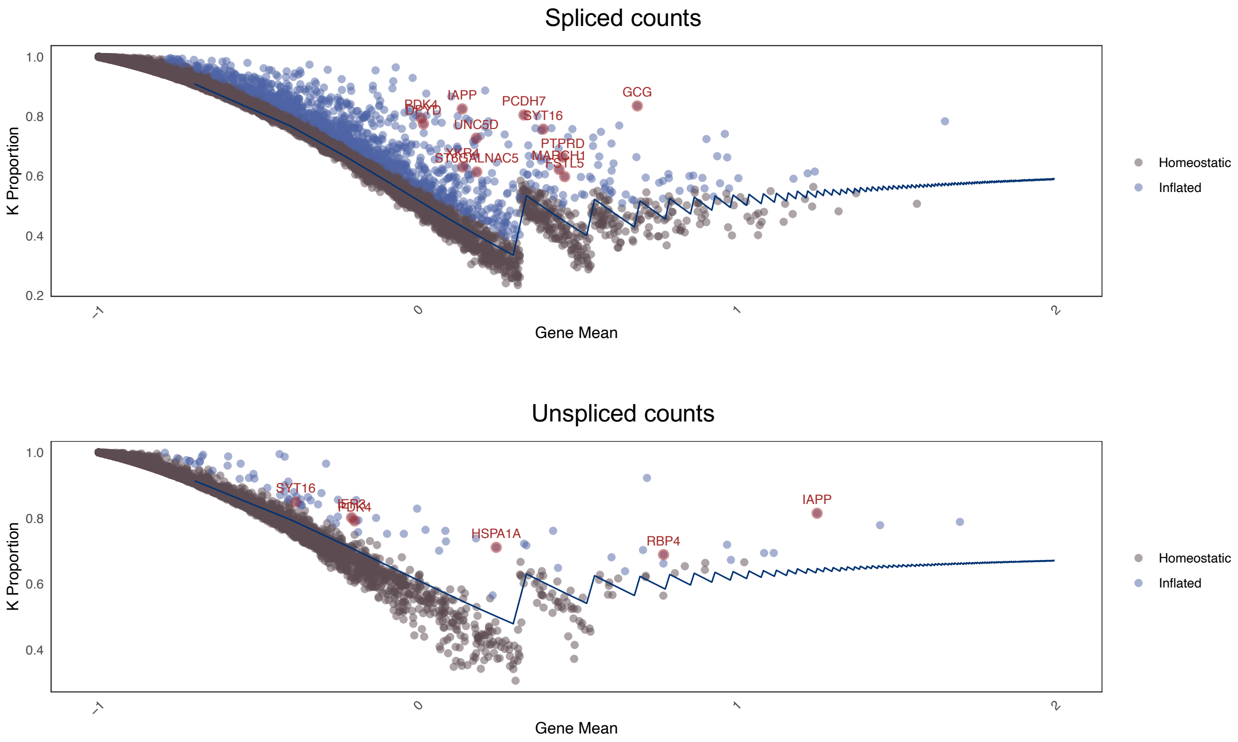


**Figure S3. Gene expression stability analysis in beta cells from the islet of one donor. We assessed the stability Z-index for both spliced and unspliced counts. Intriguingly, the k-proportions for these counts each adhere to a distinct group of negative binomial distributions, both maintaining the same dispersion level.**

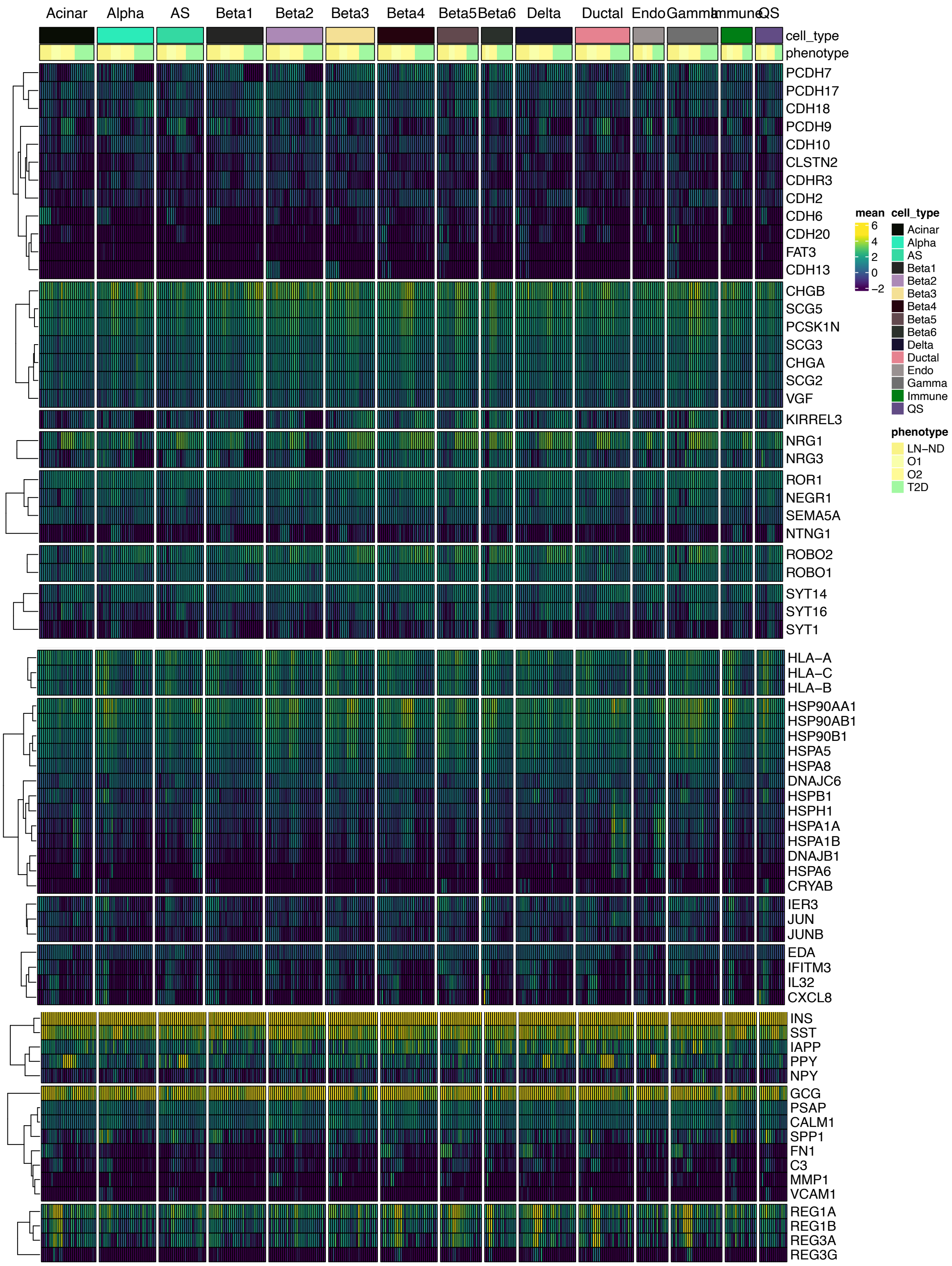


**Figure S4. Heatmap displays the mean normalized counts for each donor cross different cell types. The difference of regulatory activities revealed by stability analysis cannot be revealed by mean-based approaches.**

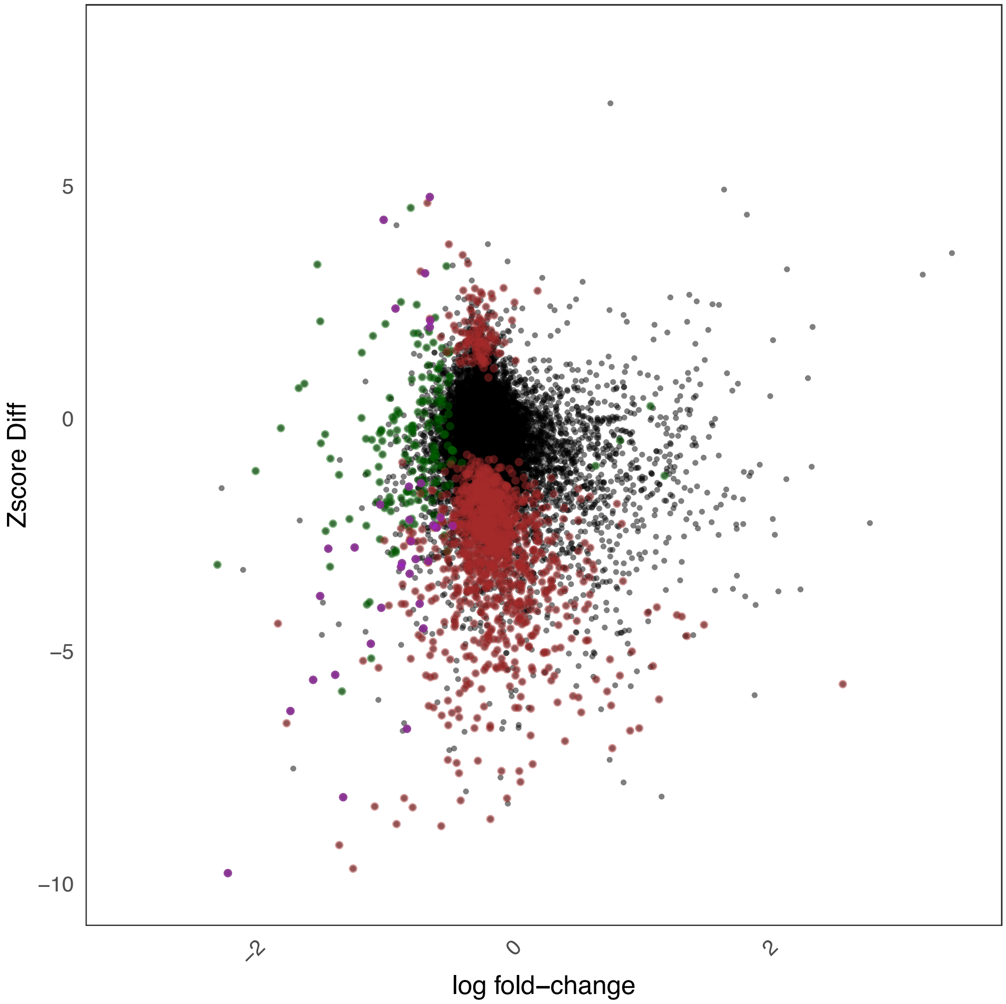


**Figure S5. Differential expression analysis using gene mean for T2D vs. ND, and compared log fold-change vs. differences in Z-index. Differential stability hits shown in red, differential expression hits shown in green and double hits shown in purple.**

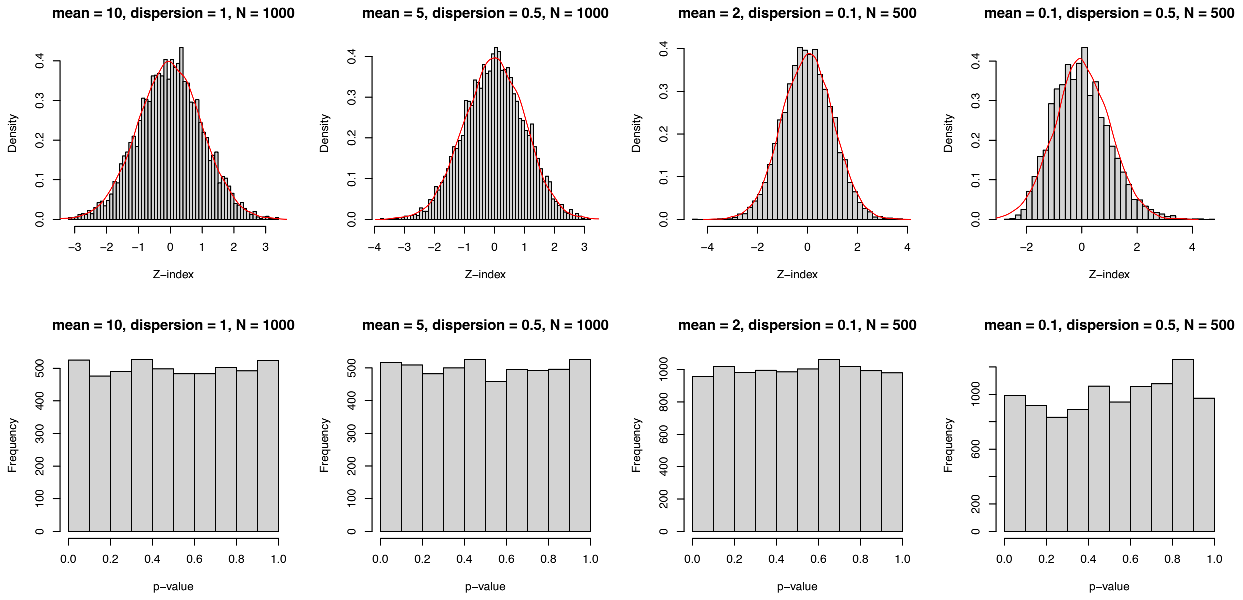


**Figure S6. Simulation studies show asymptotic normality of k-inflation test. Z-indexes and p-values are calculated under the null hypothesis.**

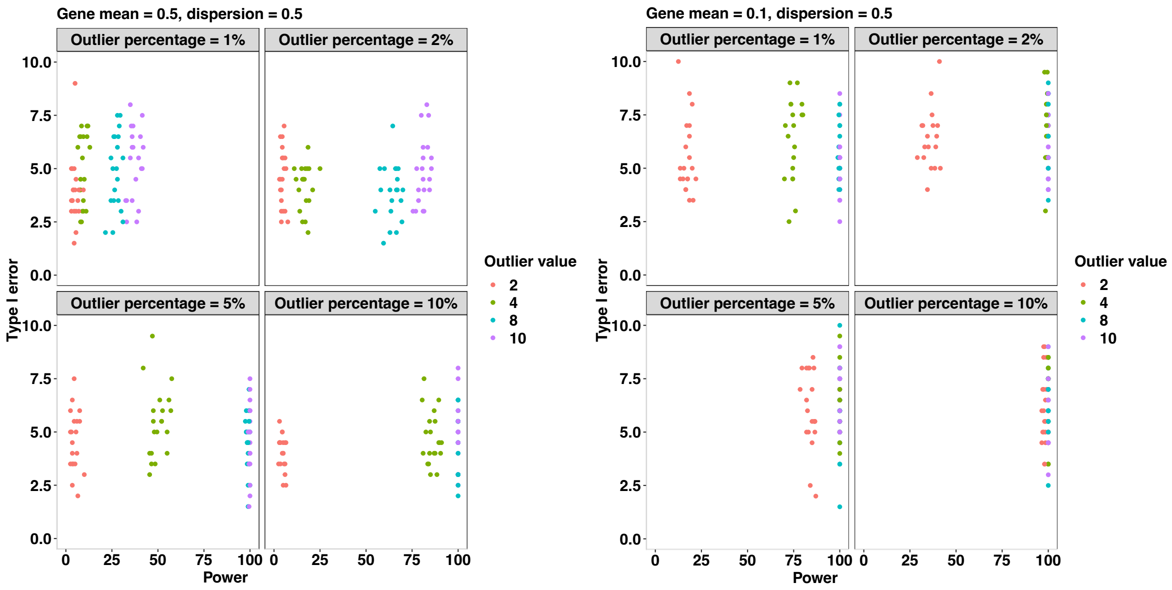


**Figure S7. Simulation studies show the power and type I error control of proposed tests, under various scenario.**

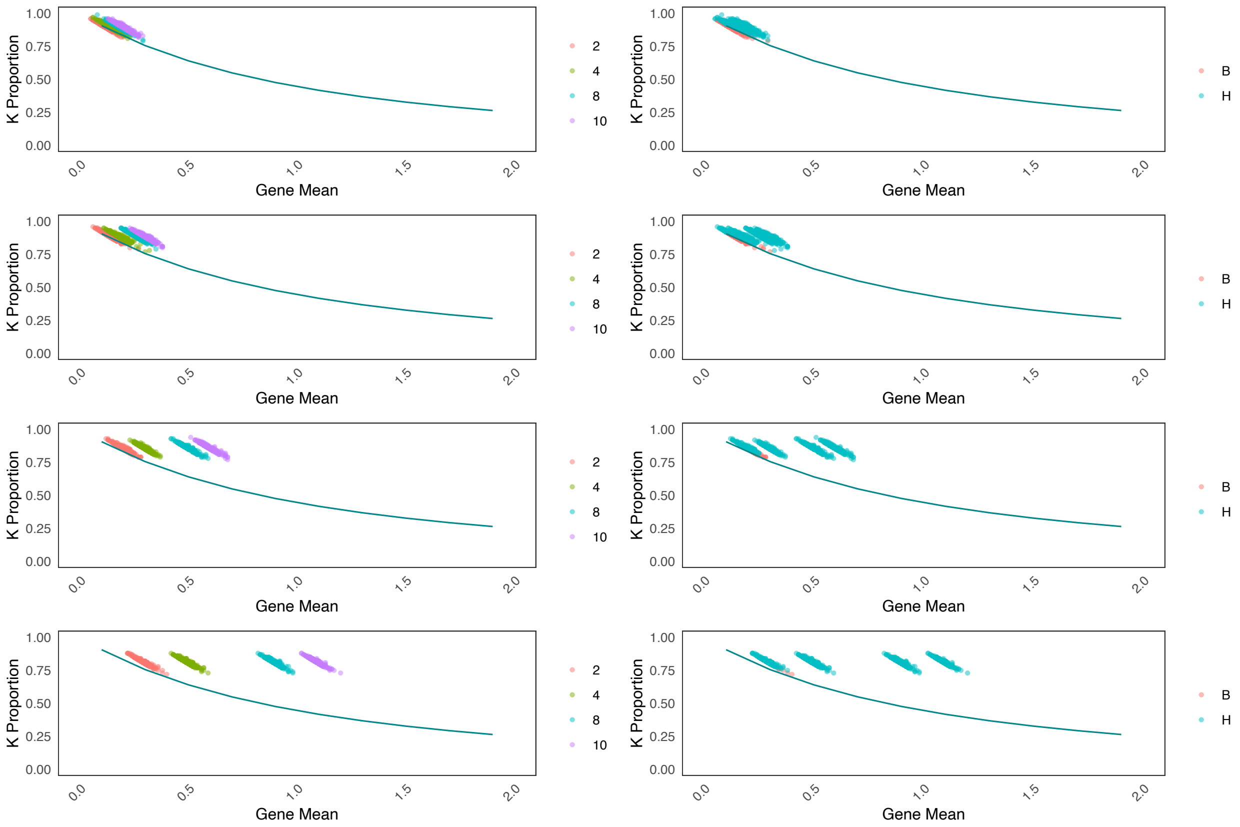


**Figure S8. Waveplots for simulation studies in Figure S7, showing the simulated inflated genes (left) and their detection using the proposed test (right). Successfully identified hits are colored in green while missed ones colored in red.**
